# Regulatory mechanism of heme-regulated inhibitor through autophosphorylation-driven activation and heme-induced deactivation

**DOI:** 10.1101/2025.10.07.681048

**Authors:** Tomomasa Oka, Hisashi Yoshida, Kenji Mizutani, Eiji Obayashi, Sam-Yong Park, Kenji Iwasaki

**Affiliations:** Degree Programs in Pure and Applied Sciences, Graduate School of Science and Technology, University of Tsukuba, 1-1-1 Tennodai, Tsukuba, Ibaraki 305-8571, Japan, University of Tsukuba, 1-1-1 Tennodai, Tsukuba, Ibaraki 305-8577, Japan; Life Science Center for Survival Dynamics, Tsukuba Advanced Research Alliance (TARA), University of Tsukuba, 1-1-1 Tennodai, Tsukuba, Ibaraki 305-8577, Japan; Graduate School of Medical Life Science, Yokohama City University, 1-7-29 Suehiro-cho, Tsurumi-ku, Yokohama 230-0045, Japan; Department of Biochemistry, Shimane University School of Medicine, 89-1 Enya-cho, Izumo 693-8501, Japan; Center for Quantum and Information Life Sciences, University of Tsukuba, 1-1-1 Tennodai, Tsukuba, Ibaraki 305-8577, Japan

**Keywords:** eIF2α, heme protein, HRI, ISR, translation

## Abstract

Heme-regulated inhibitor (HRI) is a key modulator of hemoglobin synthesis, exerting such effects by sensing intracellular heme levels. Under heme-deficient conditions, HRI dissociates from heme, becomes activated, and phosphorylates the translation initiation factor eIF2α. However, the precise regulatory mechanisms governing HRI activation remain incompletely understood. In this study, we delineate part of the regulatory mechanism involving autophosphorylation-dependent activation and heme-mediated deactivation of HRI. HRI formed a dimer in solution through its N-terminal domain, irrespective of its phosphorylation state. However, a mutant with deletion of the N-terminal domain retained autophosphorylation activity, indicating that N-terminal domain-mediated dimerization is not essential for activation. Phosphorylated HRI formed a stable complex with eIF2α, whereas the dephosphorylated form failed to bind, indicating that autophosphorylation is required for eIF2α recognition. Biochemical analyses, together with modeling based on the predicted structure, revealed that the phosphate groups of Thr488 and Thr493 interact with adjacent basic residues. These interactions enhance eIF2α recognition and phosphorylation. Additionally, heme-induced deactivation selectively targeted the dephosphorylated kinase domain and suppressed autophosphorylation. These findings provide multiple mechanistic insights into regulation of the activity of HRI, highlighting distinct roles for autophosphorylation, structural elements, and heme responsiveness in controlling its function.

---

Eukaryotic cells employ multiple regulatory mechanisms to ensure tight control over translation, a process essential for synthesizing proteins from mRNA. One such mechanism is the integrated stress response (ISR), which is triggered by specific cellular stressors. Activation of the ISR leads to a global reduction in translation, thereby preventing the accumulation of misfolded proteins (*1*). Initiation of the ISR is regulated by a group of kinases that specifically phosphorylate the α subunit of eukaryotic translation initiation factor 2 (eIF2α). To date, four types of eIF2α kinases have been identified: protein kinase R (PKR), PKR-like endoplasmic reticulum kinase (PERK), general control nondepressible 2 (GCN2), and heme-regulated inhibitor (HRI). Each kinase responds to distinct stressors, such as viral infection, endoplasmic reticulum stress, amino acid deprivation, and heme deficiency. Upon the perception of stress signals, these kinases are activated and phosphorylate Ser51 of eIF2α, preventing nucleotide exchange on eIF2γ and halting translation (*2,3*). Therefore, sensing stress and activating eIF2α kinases are critical steps in ISR initiation; however, the activation mechanism differs for each eIF2α kinase (*4,5*).

HRI senses heme deficiency and inhibits translation. It is predominantly expressed in reticulocytes and red blood cells, where it regulates globin synthesis in response to heme availability (*6*). HRI is composed of two domains, the N-terminal domain (residues 1–144) and the kinase domain (residues 160–630), including a kinase insert at residues 235–263, as shown in Fig. 1. HRI activation is thought to occur through the following steps. Under normal conditions, HRI remains in a dephosphorylated state (off state) due to its association with heme, mediated by the two domains (*7,8*). HRI dissociates from heme under heme-deficient conditions, depending on the heme concentration. Subsequently, free HRI autophosphorylates and becomes active, phosphorylating Ser51 of eIF2α and thereby stopping translation globally. Recently, novel functions of HRI beyond the modulation of hemoglobin translation have become evident (*6,9,10*). For instance, it has been reported that HRI is expressed in neurons and halts protein translation in response to the accumulation of denatured proteins (*11,12*). While HRI is primarily known for heme sensing in erythroid cells, emerging evidence suggests broader functions in other tissues, such as neurons.

**Fig. 1.**
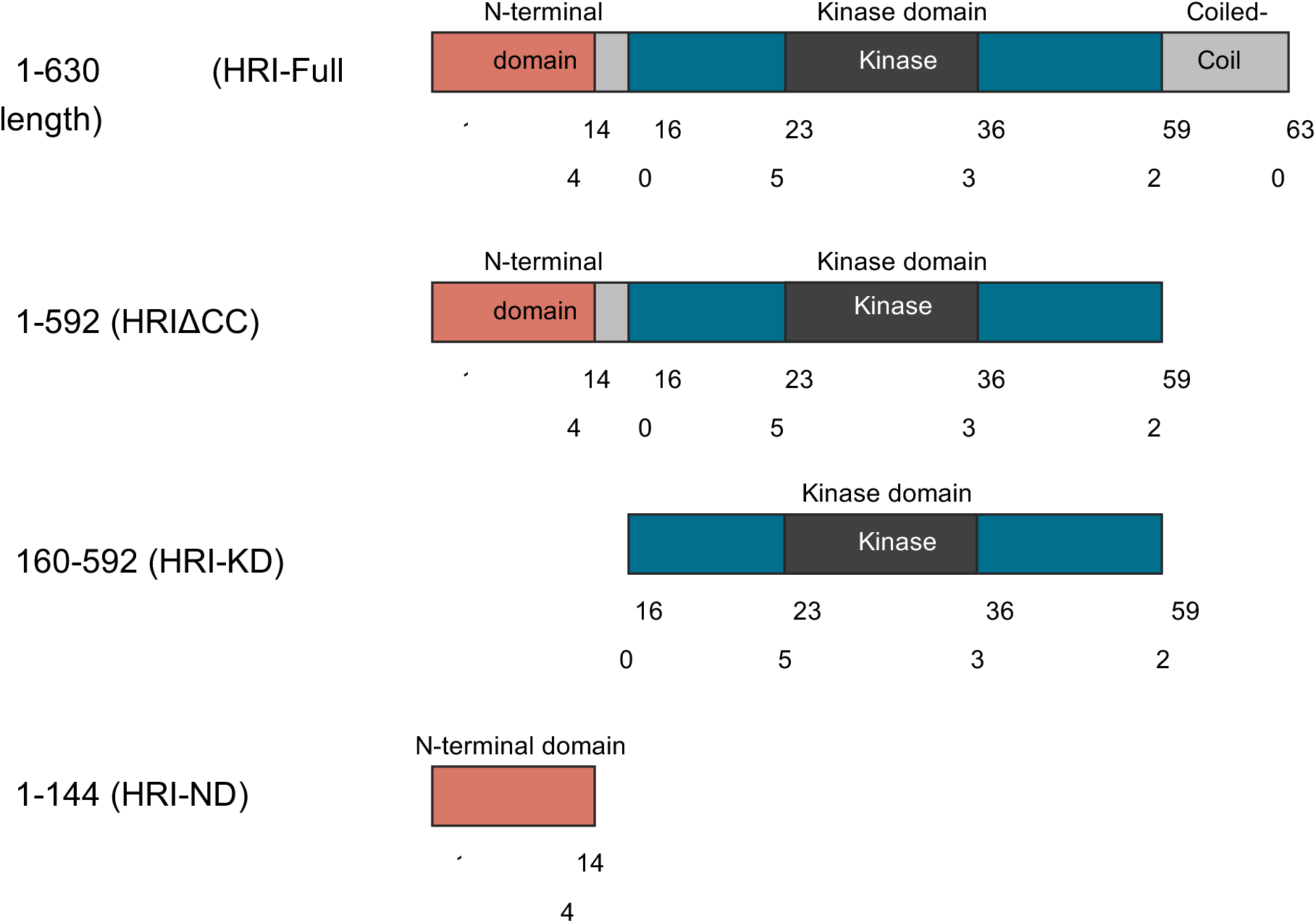
Domain maps of HRI and its four engineered deletion constructs.

Several heme-binding proteins, such as cytochrome P450, function by forming active centers via heme coordination. Interestingly, some of these proteins act as heme sensors through reversible heme binding and are involved in diverse biological processes including transcription, translation, and heme biosynthesis (*13*). For instance, PefR regulates the transcription of heme efflux-related genes in response to intracellular heme concentrations (*14*). Many heme-sensor proteins possess a conserved Cys-Pro (CP) motif that serves as an axial ligand for heme (*15–17*). Igarashi et al. proposed that His119/120 in the N-terminal domain and the CP motif (C410, P411) in the kinase domain of *Mus musculus* HRI (mHRI) coordinate heme binding (*7*). Additional residues, including His78 and His123, have also been implicated in heme coordination (*18*). These findings support the view that HRI functions as a heme sensor protein.

Despite extensive studies, the molecular mechanisms underlying HRI activation and heme-induced deactivation remain incompletely understood. As many as 33 autophosphorylation sites have been identified in mHRI, spanning a wide range of its sequence (*19,20*). Among them, phosphorylation at Thr485 and Thr490 has been suggested to play a key role in catalytic activation (*19*). As with many kinases, self-multimerization often triggers autophosphorylation (*21*); however, it remains unclear how HRI autophosphorylates multiple sites and how this process regulates its catalytic activity. AlphaFold structural prediction suggests that His119/120 and C410–P411, both proposed as potential axial ligands for heme, are located far apart, raising questions about the precise heme-binding conformation (22). Although several heme-coordinating residues have been proposed, the mode of heme binding and the mechanism by which heme suppresses HRI activity remain elusive.

HRI has recently emerged as a potential therapeutic target for sickle cell disease, which results from abnormal hemoglobin polymerization and leads to severe anemia. One promising therapeutic approach involves upregulating fetal hemoglobin (HbF) expression. In HRI- knockdown erythroid cells, HbF production increased (*23,24*), suggesting that modulation of HRI activity could provide clinical benefit. Therefore, elucidating the mechanism of HRI activation is not only of biological interest but may also inform therapeutic strategies. In this study, we aimed to clarify how autophosphorylation and heme binding govern HRI activity. We present evidence for the involvement of multimerization in autophosphorylation, its impact on eIF2α binding, and the respective roles of the N- and C-terminal domains in heme coordination.

## Experimental procedures

### Cloning and mutagenesis

The genes encoding human HRI (Gene ID: 27102) and human eIF2α (Gene ID: 1965) were amplified by polymerase chain reaction (PCR) using a cDNA library derived from HEK293T cells. The hHRI deletion lacking the C-terminal coiled-coil (1–592, HRIΔCC), the hHRI kinase domain (160–592, HRI-KD), the hHRI N-terminal domain (1–144, HRI-ND), and the human eIF2α gene (2–183) were amplified by PCR from full-length cDNA as a template. The products were digested with BamHI and NotI, and ligated into a suitably cut, modified pET28b vector in which an SD sequence, an initial ATG, a hexahistidine tag, and a tobacco etch virus (TEV) protease cleavage site had been cloned between the XbaI and BamHI restriction sites immediately upstream of the protein-coding region. Site-directed mutagenesis was carried out by PCR to generate HRI-KD (C411S), HRIΔCC (T488/493A), and HRIΔCC lacking the kinase insert (residues 234–363).

### Protein expression and purification

All constructs were expressed in BL21 (DE3) or Rosetta (DE3) strains. The transformed cells were cultured at 37°C until absorbance of 0.6–1.0 at 600 nm was reached. Subsequently, isopropyl-β-D-thiogalactoside was added to a final concentration of 0.5 mM, and the cells were cultured with shaking at 18°C overnight. After centrifugation (6000g, 10 min), the cells were resuspended in Ni-NTA binding buffer [20 mM Tris (pH 8.0), 500 mM NaCl, 500 mM urea, 20 mM imidazole, 5% glycerol, and 2 mM 2-mercaptoethanol] and lysed by sonication. After centrifugation (20,000 g, 30 min), the supernatant was loaded onto a Ni-NTA agarose column (QIAGEN, Hilden, Germany) equilibrated with Ni-NTA binding buffer. After incubation for 1 h, the protein was eluted with Ni-NTA elution buffer [20 mM Tris-HCl (pH 8.0), 500 mM NaCl, 500 mM urea, 500 mM imidazole, 5% glycerol, and 2 mM 2-mercaptoethanol]. His-TEV protease was added to the elution fraction, and it was dialyzed against Ni-NTA binding buffer at 4°C overnight. The solution was reloaded onto the same Ni-NTA agarose and incubated at 4°C for 1 h, after which the flow-through was collected. The samples were dialyzed against Buffer A (20 mM Tris-HCl [pH 8.0], 100 mM NaCl, and 2 mM 2-mercaptoethanol). HRI was loaded onto a Hitrap Q column (Cytiva, Marlborough, MA, USA) and eluted with a linear gradient of 100–1000 mM NaCl. The peak fractions were collected and loaded onto a HiLoad 16/600 column (Cytiva) equilibrated with buffer A. To produce dephosphorylated HRI, cells expressing HRI were co-sonicated with cells expressing λ-phosphatase (gene ID: 2703537) and incubated for 1 h at room temperature, after which the samples were purified using the same procedure as for phosphorylated HRI. Following affinity purification, eIF2α was loaded onto a HiLoad 16/600 column equilibrated with 20 mM Tris-HCl (pH 8.0), 500 mM NaCl, and 2 mM 2- mercaptoethanol. All samples were concentrated using Amicon Ultras (Millipore, Billerica, MA, USA) and stored at −80°C until use.

### Kinase activity assay

Kinase activity was measured based on a previous report with minor modifications (*7*). HRI and eIF2α were mixed at final concentrations of 0.5 μM and 5 μM, respectively, in a reaction buffer containing 20 mM Tris-HCl (pH 7.7), 60 mM KCl, and 2 mM magnesium acetate. The mixture was incubated at 15°C or 16°C for 15 min, and the reaction was initiated by adding ATP to a final concentration of 350 μM. The reaction was terminated at the specified time points by adding SDS loading dye. For assays with hemin, HRI (final concentration: 2 μM) was incubated overnight at 4°C with hemin at a final concentration of 0, 10, 20, or 40 μM in a buffer containing 20 mM Tris-HCl (pH 7.7), 60 mM KCl, 2 mM magnesium acetate, and 5% (v/v) DMSO. HRI and eIF2α were mixed as described above in a reaction buffer containing 20 mM Tris-HCl (pH 7.7), 60 mM KCl, 2 mM magnesium acetate, and hemin dissolved in 5% v/v DMSO at a final concentration of 0, 10, 20, or 40 μM. The reaction was initiated and terminated as described above. The samples were denatured at 95°C for 5 min and loaded onto 12% (w/v) SDS-PAGE gel containing 100 μM Phos-tag acrylamide (NARD Institute, Kobe, Japan) and 100 mM manganese chloride.

### Qualitative interaction analysis by size exclusion chromatography (SEC)

HRI and eIF2α were mixed at a molar ratio of 1:1.2 or 1:1.5 in a buffer containing 20 mM Tris-HCl (pH 8.0), 100 mM NaCl, and either 1 mM DTT or 2 mM 2-mercaptoethanol. After 1 h of incubation at 4°C, the mixtures were loaded onto a HiLoad 16/600 column, and the peak fractions were analyzed by SDS-PAGE.

### Biolayer interferometry (BLI) assays

Quantitative analysis of the interaction between HRIΔCC or HRIΔCC (T488/493A) and eIF2α (2–183) was performed using BLI assays. HRI samples were expressed and purified as described above, while eIF2α (2–183) was prepared in the same manner but without His-tag cleavage. BLI assays were conducted using the Octet N1 (Sartorius, Göttingen, Germany) equipped with Octet® HIS1K Biosensors, with shaking at 2200 rpm and room temperature for all assays. Before the assays, the samples were dialyzed in 20 mM Tris-HCl (pH 8.0), 100 mM NaCl, and 1 mM TCEP. After dialysis, BSA was added to the samples to a final concentration of 0.1% (w/v). The assay buffer was identical to the buffer used to dilute the samples. BLI was performed as follows: (1) HIS2K biosensors were immersed in the assay buffer for 30 s; (2) the biosensors were immersed in a 50 ng/μL His-tagged eIF2α solution for 120 s to load eIF2α onto the sensors; (3) the biosensors were immersed in the assay buffer for 30 s for washing; (4) the biosensors were immersed in HRI solutions (50, 10, 5, and 1 nM) for 120 s; and (5) the biosensors were immersed in the assay buffer for 600 s to dissociate HRI from the sensors. The experimental data were analyzed using a global fitting model (1:1) with at least four curves per replicate to calculate the dissociation constant (*K*_d_).

### Analytical ultracentrifugation (AUC) and mass photometry

The molecular weight (MW) and self-association of HRI were examined using a Beckman Optima XL-I analytical ultracentrifuge (Beckman Coulter, Fullerton, CA, USA). The experiments were performed at 20 °C, and the absorbance at 280 nm was measured. Before the experiment, the sample was dialyzed in 20 mM Tris-HCl (pH 8.0), 100 mM NaCl, and 1 mM TCEP. Following 2 h of thermal equilibration, sedimentation velocity data were obtained every 10 min for 17 h at 40,000 rpm. The obtained data were subsequently analyzed using a Cs distribution model in the program SEDFIT (*25*). The frictional ratio was allowed to float during fitting. The partial specific volume, solvent density, and viscosity were calculated by SEDNTERP (*26*). The estimation of MW was also confirmed by mass photometry using a Refeyn, Two MP (Refeyn Ltd., Oxford, UK) (*27*). Purified dephosphorylated HRIΔCC was dialyzed against 20 mM Tris-HCl (pH 8.0), 100 mM NaCl, and 2 mM TCEP, and its MW was measured following the manufacturer’s procedure.

### Hemin titration assay

Dephosphorylated HRI was dissolved in 20 mM Tris-HCl (pH 8.0), 200 mM NaCl, and 5% DMSO at a concentration of 10 μM, and 400 μl of the solution was injected into a cuvette. Hemin was dissolved in the same buffer at a concentration of 200 μM. Hemin titration was performed by adding 2 μL aliquots of the hemin solution. The change in absorption was measured using a NanoDrop™ (Thermo Fisher Scientific, Cleveland, OH, USA).

### Generation of predicted structural model

The structural model of hHRI-KD, with Thr488 and Thr493 either phosphorylated or unphosphorylated, was predicted using AlphaFold 3 (*28*). The amino acid sequence used for model prediction was obtained from UniProt with the following accession number: hHRI: Q9BQI3.

## Results

### N-terminal domain-dependent dimerization

We first attempted to express and purify full-length human HRI, a C-terminal coiled-coil deletion mutant (residues 1–592; HRIΔCC), the isolated N-terminal domain (residues 1–144; HRI-ND), and the isolated kinase domain (residues 160–592; HRI-KD) (Fig. 1). Previous studies have reported that HRI expressed in *E. coli* is extensively phosphorylated (19). Therefore, we employed an *E. coli* expression system for all constructs. HRIΔCC, HRI-ND, and HRI-KD were obtained in soluble form whereas full-length HRI was localized exclusively to the insoluble fraction. SEC profiles of HRIΔCC, HRI-KD, and HRI-ND are shown in Fig. 2A. The elution volumes of HRIΔCC, HRI-KD, and HRI-ND were 63.8, 75.7, and 71.6 mL. The elution volume of HRI-KD was greater than that of HRI-ND, although their theoretical MWs are 49 and 16, respectively. To assess the oligomeric states of the constructs, we performed sedimentation velocity analysis. The sedimentation coefficients (Sw) of HRIΔCC and HRI-KD were 7.4 and 3.1 (Fig. 2B), corresponding to the estimated MWs of 127 for HRIΔCC and 58 for HRI-KD, respectively. While the estimated MW of HRI-KD was consistent with the theoretical value (49), that of HRIΔCC was approximately twice the theoretical MW (69), indicating that HRIΔCC forms a dimer and HRI-KD is monomeric. The Sw and estimated MW of HRI-ND were 3.7 and 39, respectively. The estimated MW was approximately twice its theoretical MW (16), demonstrating that the N-terminal domain forms a dimer. These data collectively indicate that both HRIΔCC and HRI-ND exist as dimers, whereas HRI-KD remains monomeric.

**Fig. 2.**
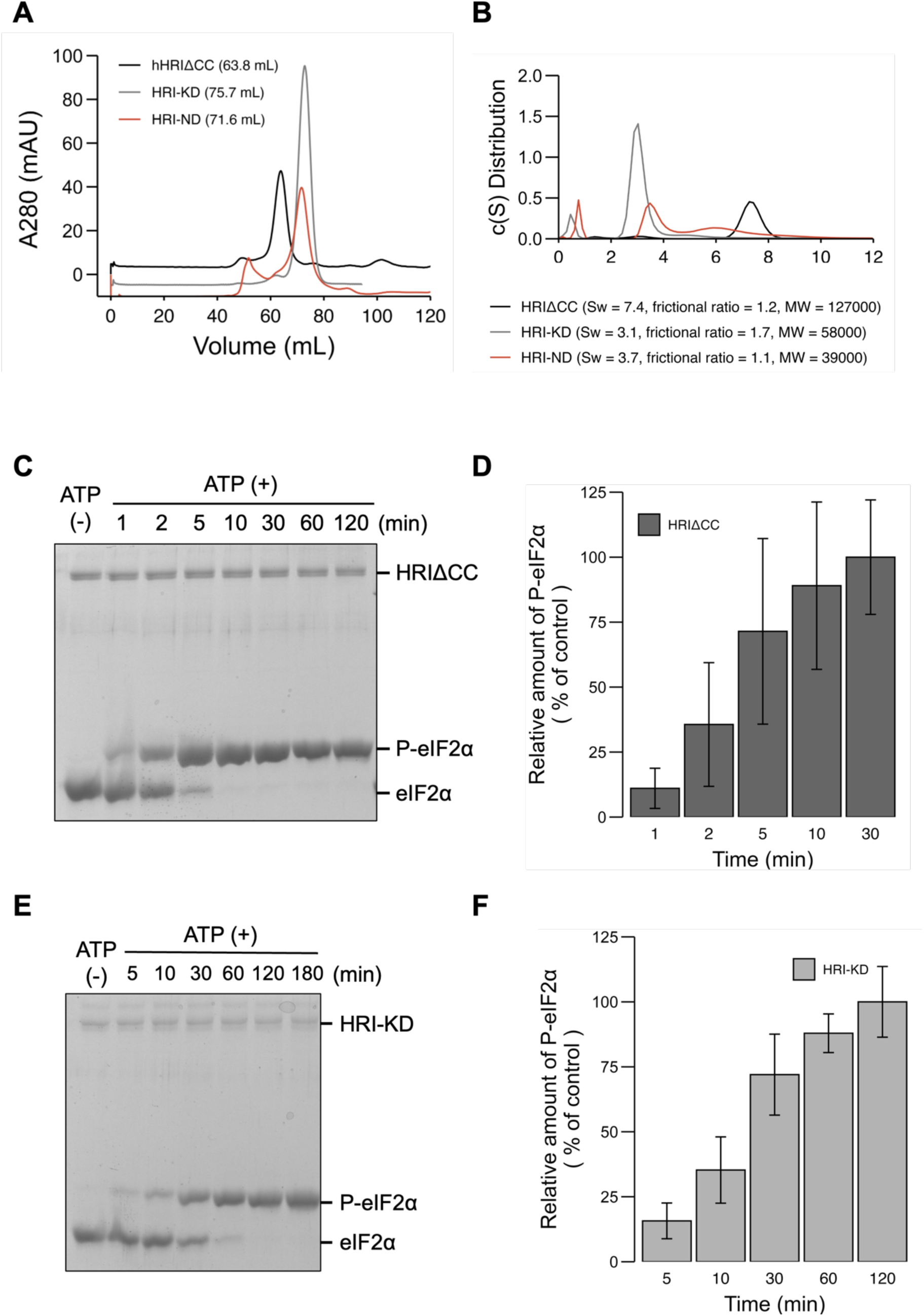
Effects of N-terminal domain on dimerization and kinase activity in phosphorylated HRI. (*A*) SEC chromatograms of HRIΔCC, HRI-KD, and HRI-ND, shown in black, gray, and red, respectively. **(*B*)** Sedimentation velocity analysis of HRIΔCC, HRI-KD, and HRI-ND. Colors correspond to those used in **(*A*)**. **(*C*)** Kinase activity of HRIΔCC toward eIF2α analyzed using Phos-tag SDS-PAGE. ***(D)*** Quantification of HRIΔCC kinase activity based on the data shown in **(*C*)**. The level of phosphorylated eIF2α at 30 min was set as a control. Error bars represent the standard error. Mean values were obtained from three independent experiments. **(*E*)** Kinase activity of HRI-KD. **(*F*)** Quantification of HRI-KD kinase activity. Data were obtained from the gel shown in **(*E*)**. The level of phosphorylated eIF2α at 120 min was set as the control. Error bars represent the standard error. Mean values were obtained from three independent experiments.

To assess the kinase activity of purified HRI and the contribution of the N-terminal domain to this activity, we performed phosphorylation assays using phos-tag acrylamide. Phos-tag acrylamide retards electrophoretic mobility by trapping phosphorylated proteins, with the degree of retardation proportional to the number of phosphate groups present. A phosphorylated eIF2α band was detected in the presence of ATP (Fig. 2C). HRIΔCC phosphorylated 1 nmol of eIF2α in a time-dependent manner, completing the reaction within 30 min (Fig. 2C and D). While HRI-KD demonstrated activity comparable to that of HRIΔCC, it required 120 min to achieve complete phosphorylation of eIF2α under identical conditions (Fig. 2E and F). These results demonstrate that purified HRI is catalytically active and that the kinase activity of HRIΔCC exceeds that of HRI-KD.

### The role of N-terminal domain in autophosphorylation

To examine how phosphorylation status influences N-terminal domain-dependent dimerization and kinase activity, we prepared dephosphorylated HRI by co-sonicating *E. coli* expressing HRI and λ-phosphatase, followed by purification as for the phosphorylated form. The SEC chromatograms of dephosphorylated HRI protein are shown in Fig. 3A. Dephosphorylated HRIΔCC [(-P)-HRIΔCC] and dephosphorylated HRI-KD [(-P)-HRI-KD] eluted as single peaks at 68.5 mL and 79.3 mL, respectively. Compared with the phosphorylated forms, both proteins exhibited an ∼5 mL increase in elution volume (Figs. 2A and 3A), suggesting either a decrease in MW or a conformational change upon dephosphorylation. Sedimentation velocity analysis showed that (-P)-HRIΔCC had an Sw of 5.9 and an estimated MW of 107 (Fig. 3B), ∼20 lower than that of the phosphorylated form. The frictional ratio was 1.2 for phosphorylated HRIΔCC and 1.4 for the dephosphorylated form (Fig. 2B and 3B). Because molecular shape can influence the results of sedimentation velocity analysis, we performed mass photometry analysis. This analysis yielded an MW of 115 for (-P)-HRIΔCC, similar to the phosphorylated form (Fig. S1). In addition, (-P)-HRI-KD exhibited an Sw of 3.0 and an estimated MW of 53, both comparable to those of the phosphorylated form. These results indicate that N-terminal domain-dependent dimerization of HRIΔCC is maintained after dephosphorylation, although the molecular shapes of the phosphorylated and dephosphorylated forms may differ.

**Fig. 3.**
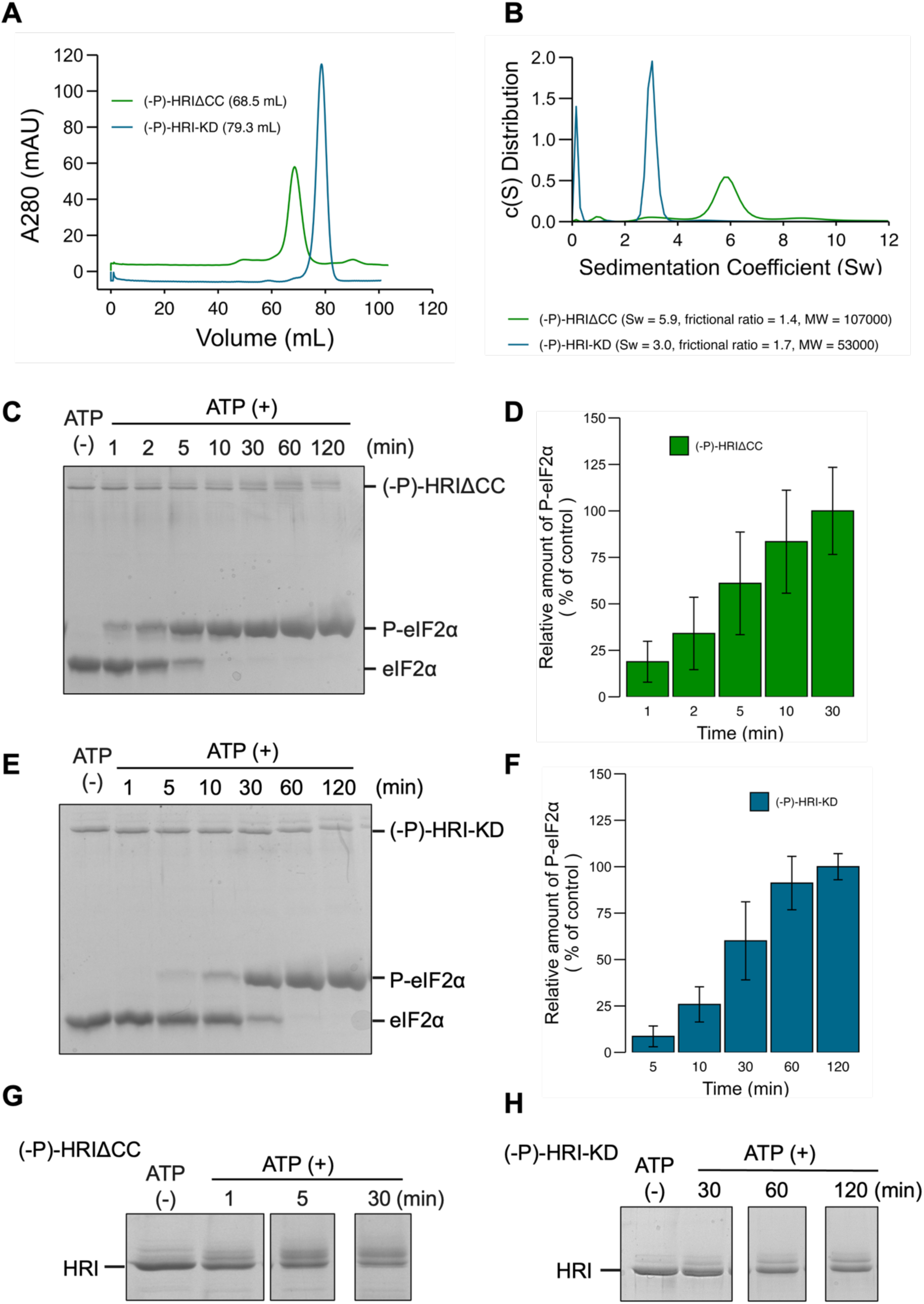
Effects of N-terminal domain on dimerization and kinase activity in dephosphorylated HRI. (*A*) SEC chromatograms showing the peak volumes of (-P)-HRIΔCC and (-P)-HRI-KD. Green and cyan represent (-P)-HRIΔCC and (-P)-HRI-KD, respectively. **(*B*)** Sedimentation velocity analysis of (-P)-HRIΔCC and (-P)-HRI-KD. The same colors are used as in **(*A*)**. **(*C*)** Kinase activity of (-P)-HRIΔCC. **(*D*)** Quantification of the kinase activity shown in **(*C*)**. The level of phosphorylated eIF2α at 30 min post-reaction was set as a control. Error bars represent standard error. Mean values were obtained from three independent experiments. **(*E*)** Kinase activity of (-P)-HRI-KD. **(*F*)** Quantification of the kinase activity shown in **(*E*)**. The level of phosphorylated eIF2α at 120 min post- reaction was set as a control. Error bars represent standard error. Mean values were obtained from three independent experiments. **(*G*)** Autophosphorylation of (-P)- HRIΔCC. **(*H*)** Autophosphorylation of (-P)-HRI-KD.

We next evaluated kinase activity to assess the role of the N-terminal domain in dephosphorylated HRI. (-P)-HRIΔCC completely phosphorylated 1 nmol of eIF2α within 30 min (Fig. 3C, D). This activity was comparable to that of phosphorylated HRIΔCC (Figs. 2D, 3D). (-P)-HRI-KD also phosphorylated 1 nmol of eIF2α, completing the reaction within 120 min, similar to the phosphorylated form (Figs. 2F and 3E, F). These results demonstrate that dephosphorylated HRI retains catalytic activity. In addition, the bands of (-P)-HRIΔCC and (-P)-HRI-KD displayed an upward mobility shift on Phos-tag acrylamide gel in a time-dependent manner (Fig. 3G, H), indicating autophosphorylation upon ATP treatment, regardless of N-terminal domain truncation. Notably, the dephosphorylated HRI bands did not fully shift upward, even when eIF2α phosphorylation was complete, suggesting that complete autophosphorylation is not required for eIF2α phosphorylation. However, it remains possible that only partially autophosphorylated molecules may still be catalytically active and produce fully phosphorylated eIF2α.

### The mechanism of HRI activation by autophosphorylation

Previous research suggests that autophosphorylation is critical for regulating HRI kinase activity (*19*). In our assay, however, dephosphorylated HRI retained kinase activity comparable to that of its phosphorylated counterpart in the presence of ATP, showing no measurable difference. We therefore examined whether phosphorylation influences the interaction of HRI with eIF2α. To address this, we compared the binding of phosphorylated and dephosphorylated HRI to eIF2α. SEC revealed that HRIΔCC mixed with eIF2α produced two peaks at 62.8 mL and 99.9 mL (Fig. 4A). SDS-PAGE analysis confirmed that HRIΔCC co-eluted with eIF2α in the 62.8 mL fraction, whereas excess eIF2α appeared at 99.9 mL (Fig. 4B). In contrast, (-P)-HRIΔCC eluted predominantly at 67.9 mL, with most eIF2α detected at 99.9 mL (Fig. 4A, C). These results indicate that phosphorylated HRI formed a stable complex with eIF2α, whereas most dephosphorylated HRI failed to form this complex. However, while autophosphorylation appears to be required for stable complex formation, phosphorylation may still proceed slowly via transient and infrequent contacts between dephosphorylated HRI and eIF2α.

**Fig. 4.**
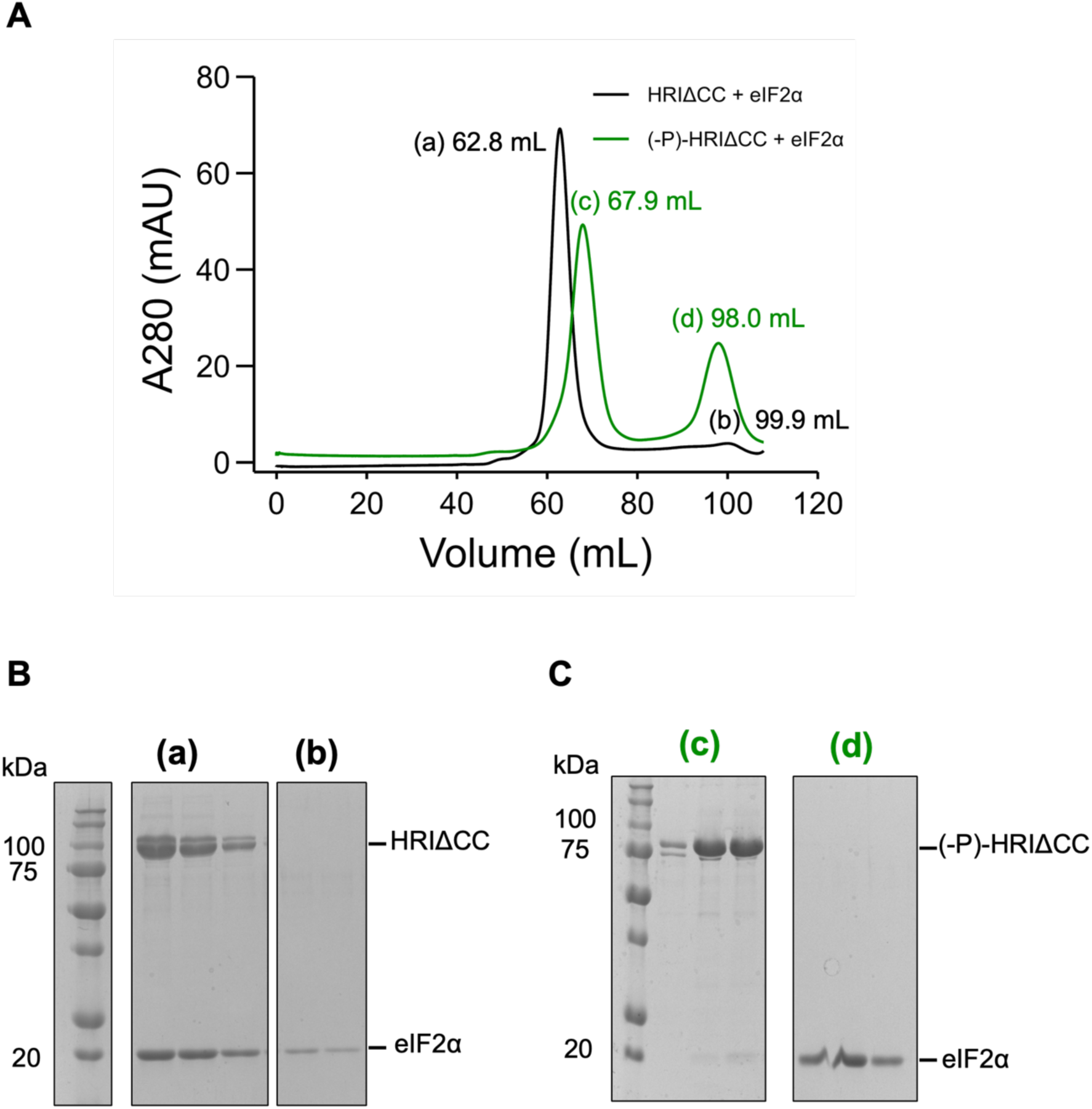
Binding capacity of phosphorylated and dephosphorylated HRI for eIF2α. (*A*) SEC chromatograms of HRIΔCC–eIF2α and (-P)-HRIΔCC–eIF2α. HRIΔCC–eIF2α and (-P)-HRIΔCC–eIF2α are shown in black and green, respectively. Detected peaks are labeled as (a) to (d). **(*B*)** SDS-PAGE analysis of fractions corresponding to peaks (a) and (b). **(*C*)** SDS-PAGE analysis of fractions corresponding to peaks (c) and (d).

Next, we sought to identify which phosphorylated residues are essential for this interaction with eIF2α. Sequence alignment between hHRI and mHRI, based on previously identified phosphorylation sites (*19*,*20*), revealed that 21 phosphorylated residues in mHRI are conserved in the hHRI kinase domain (Fig. 5). Generally, many kinases contain an activation loop (A-loop), which regulates activity through modifications such as phosphorylation or interactions with regulatory factors (*21*,*29*). This region is typically defined as spanning from the DFG (Asp-Phe-Gly) motif to the APE (Ala-Pro-Glu) motif, although some variations in this composition have been identified. In human eIF2α kinases, the DFG and SPE (Ser-Pro-Glu) motifs are conserved. In mHRI, three phosphorylation sites were identified in the A-loop, and phosphorylation at Thr485 and Thr490 proved important for kinase activity (*19*,*20*). These residues are conserved in hHRI as Thr488 and Thr493. In addition, 12 autophosphorylation sites were detected in an unstructured region known as the kinase insert loop (Fig. 5), prompting us to also investigate this region.

**Fig. 5.**
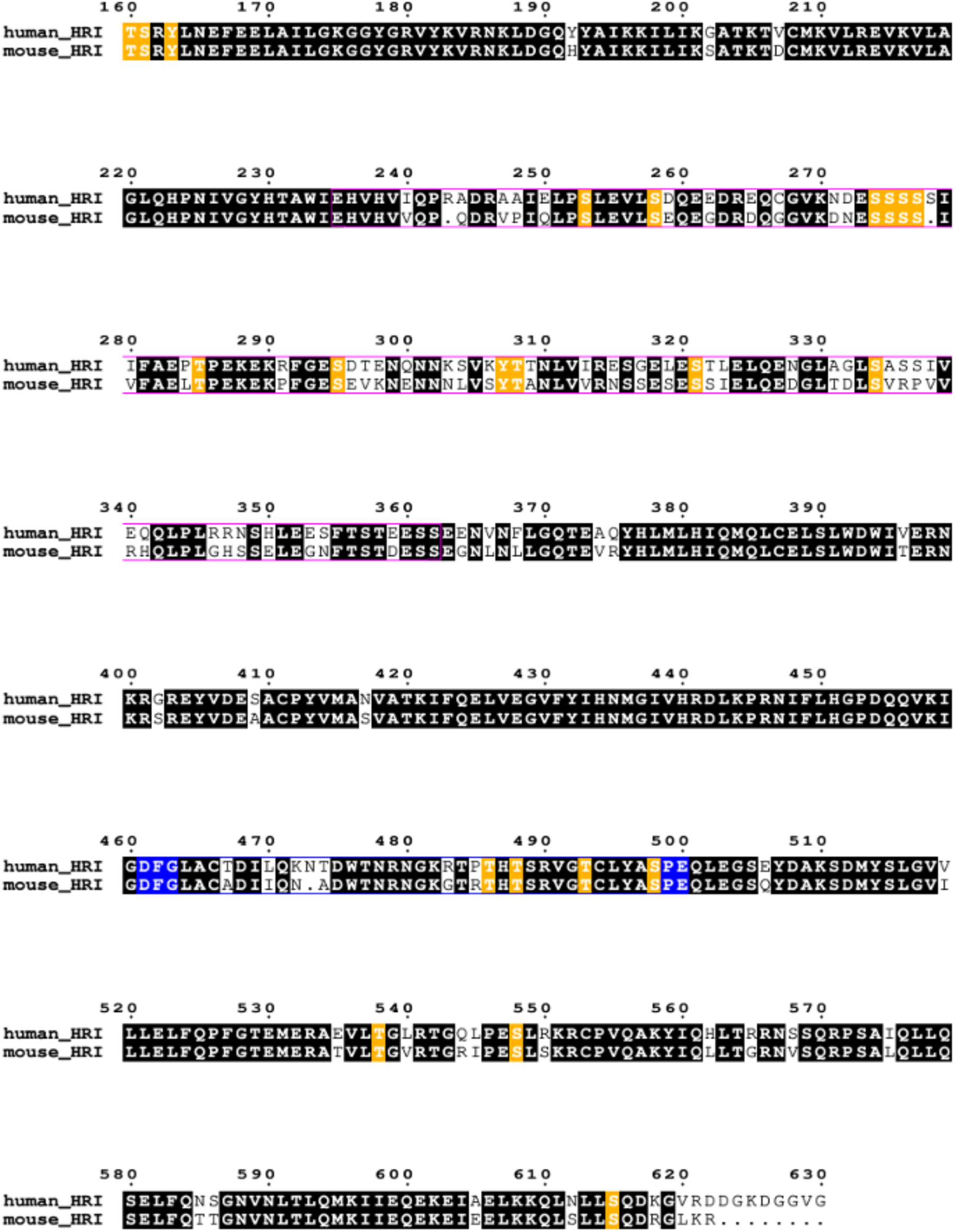
Sequence alignment of human and mouse HRI. Sequence alignment was performed using *ClustalW*, and the figure was generated using ESPript 3.0. Residues conserved between human and mouse HRI are shown in black. Amino acids identified as autophosphorylation sites in mHRI and conserved in the human sequence are highlighted in orange. The A-loop is indicated in blue, and the kinase insert loop is indicated in pink. UniProt accession numbers: human: Q9BQI3; mouse: Q9ER97.

To determine whether mutations at these residues affect kinase activity, we generated HRIΔCC mutants in which Thr488 and Thr493 were substituted with alanine. HRIΔCC (T488/493A) failed to fully phosphorylate 1 nmol of eIF2α within 30 min (Fig. 6A), whereas wild-type (WT) HRIΔCC completely phosphorylated the same amount under identical conditions (Fig. 2C). We also generated an HRIΔCC construct lacking residues 235–363, referred to as HRIΔCCΔloop (Fig. 6B). After 30 min of reaction with HRIΔCCΔloop, eIF2α remained unphosphorylated (Fig. 6C). These results indicate that the kinase activity of both HRIΔCC (T488/493A) and HRIΔCCΔloop is lower than that of WT.

**Fig. 6.**
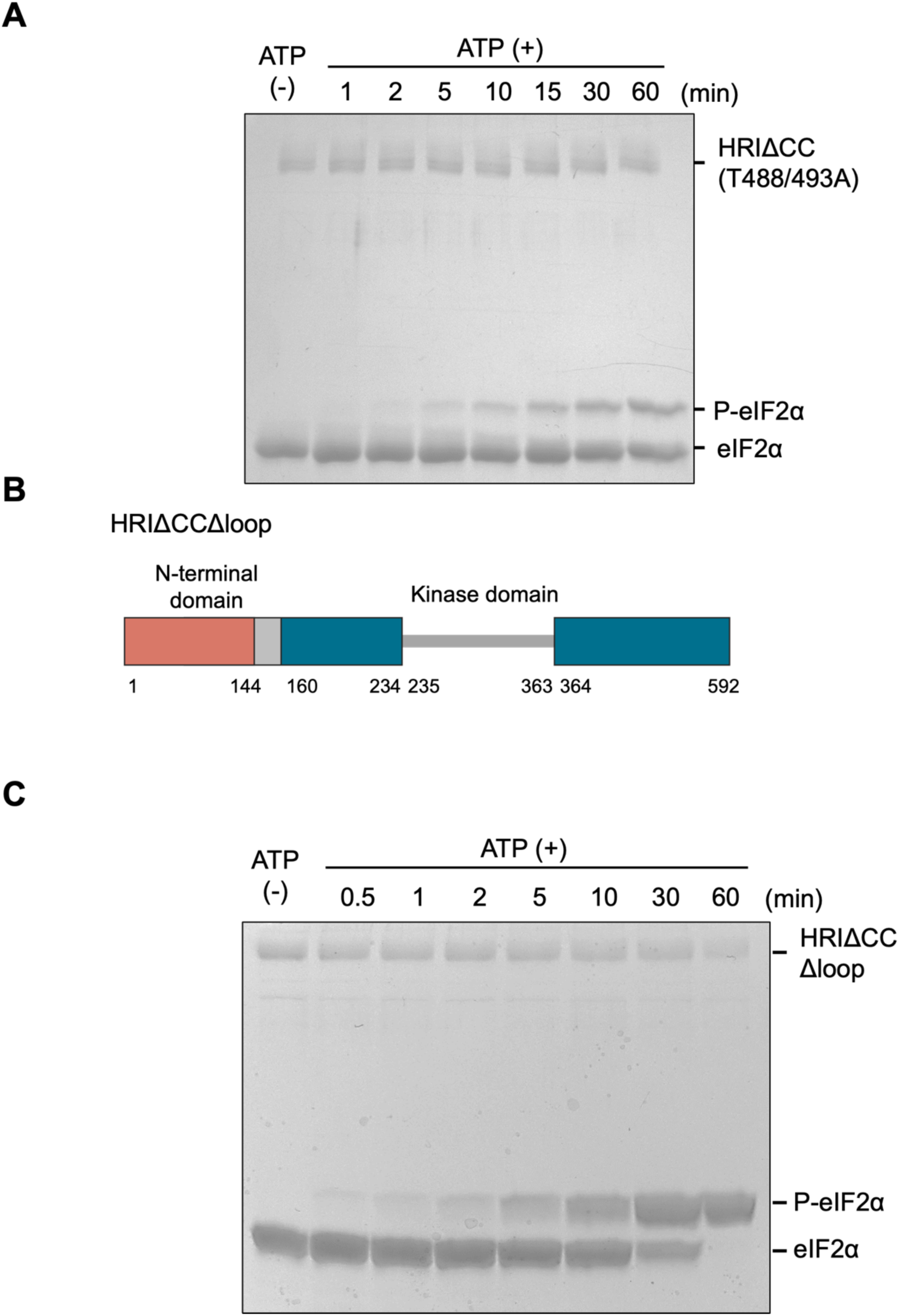
Identification of autophosphorylation sites influencing kinase activity. (*A*) Kinase activity of HRIΔCC (T488/493A). **(*B*)** Domain map of HRIΔCCΔloop. **(*C*)** Kinase activity of HRIΔCCΔloop.

To elucidate how Thr488/493 and the kinase insert loop affect kinase activity, we analyzed the interactions between the corresponding HRIΔCC mutants and eIF2α. SEC analysis of the mixture of HRIΔCC (T488/493A) and eIF2α showed a peak at 65.7 mL, demonstrating that the T488/493A mutant formed a complex with eIF2α (Fig. 7A, B). To evaluate the interaction more precisely, we performed BLI analysis. The results showed that WT HRIΔCC exhibited affinity for eIF2α, with a dissociation constant (*K*_d_) of 2.0 nM (Fig. 7C), whereas the T488/493A mutant showed a *K*_d_ of 15 nM (Fig. 7D), demonstrating reduced affinity compared with WT. When the mixture of HRIΔCCΔloop and eIF2α was applied to the SEC column, the two proteins eluted separately at 73.2 mL and 102 mL, respectively (Fig. 7E and F). This elution profile was analogous to that observed with dephosphorylated HRI and eIF2α. Thus, HRIΔCCΔloop did not form a stable complex with eIF2α.

**Fig. 7.**
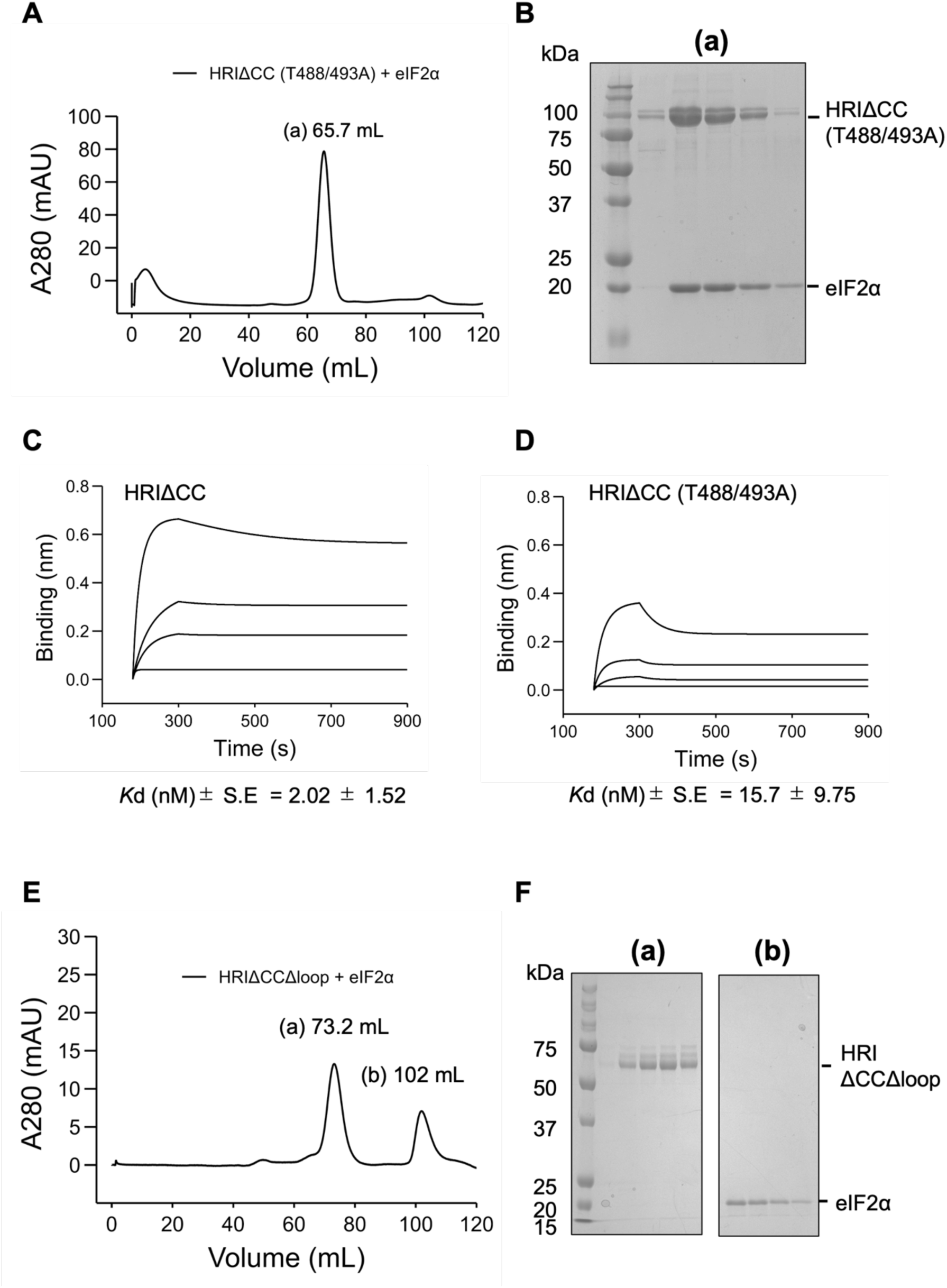
Identification of phosphorylated residues critical for interaction with eIF2α. (*A*) SEC chromatogram of the HRIΔCC (T488/493A)–eIF2α mixture. The main peak appeared at 65.7 mL (a). **(*B*)** SDS-PAGE analysis of fraction corresponding to peak (a). Binding affinities of **(*C*)** HRIΔCC and **(*D*)** HRIΔCC (T488/493A) for eIF2α. The mean dissociation constant (*K*_­­_) ± standard error was calculated from three independent experiments. **(*E*)** SEC chromatogram of HRIΔCCΔloop-eIF2α mixture. Detected peaks are labeled as (b) and (c). **(*F*)** SDS-PAGE analysis of fractions corresponding to peaks (a) and (b).

We predicted the structure of the HRI-KD to elucidate how phosphorylation at Thr488 and Thr493 affects the A-loop conformation. Fig. 8 shows the predicted structures of the dephosphorylated HRI-KD and the Thr488/493-phosphorylated HRI-KD generated using AlphaFold 3. The overall structures of the phosphorylated and dephosphorylated forms were similar, except for the kinase insert loop, which had low predictive accuracy due to its unstructured region (Fig. 8A). In contrast, the conformation of the A-loop phosphorylated at Thr488 and Thr493 differed from that of the dephosphorylated form. The phosphate group of Thr488 formed hydrogen bonds with Arg213 and Arg441, whereas the phosphate group of Thr493 interacted with Lys444 (Fig. 8B). Thr488 and Thr493 are conserved among human eIF2α kinases, despite the low overall conservation of the A-loop (Fig. 9A). In the crystal structures of the human PKR–eIF2α complex and apo mouse PERK (both members of the eIF2α kinase family), autophosphorylation of Thr446 and Thr980—corresponding to HRI Thr488—was observed (*30*, *31*). Conversely, autophosphorylation of the threonine residue corresponding to Thr493 was not observed in these structures. We compared our predicted strucure with these crystal structures to evaluate the functional role of the phosphate group at Thr488 in HRI. In the PKR structure, the phosphate group of Thr446 formed hydrogen bonds with Lys304, Arg307, and Arg413 (Fig. 9B). A similar structural arrangement was observed in PERK, where Thr980 formed interactions with Lys631, Arg634, and Arg934 (Fig. 9C). In our predicted structure, Thr488 primarily interacted with Arg213 and Arg441, but not with Lys210 (Fig. 9D). Thus, the interactions between Thr488 and surrounding basic residues observed in our predicted structure are consistent with those found in other eIF2α kinases.

**Fig. 8.**
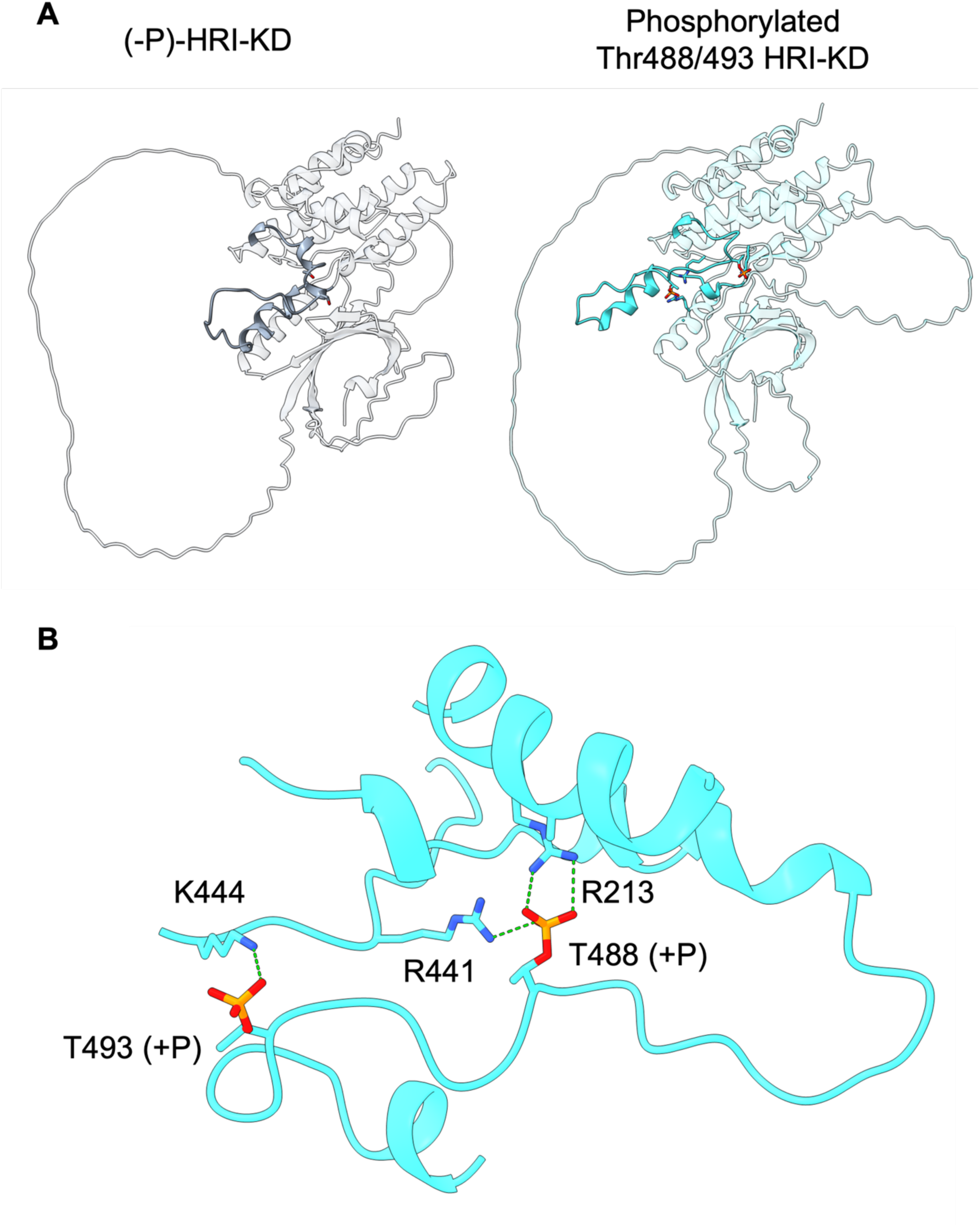
Structural differences in the A-loop between phosphorylated (Thr488/493) and dephosphorylated HRI. (*A*) Overall predicted structure of (-P)-HRI-KD (gray) and phosphorylated HRI-KD (Thr488/493; cyan). Structural models were generated using AlphaFold 3. Dark areas indicate A-loop. **(*B*)** Predicted interactions between phosphate groups and basic residues in phosphorylated HRI-KD. Hydrogen bonds are shown as green dotted lines.

**Fig. 9.**
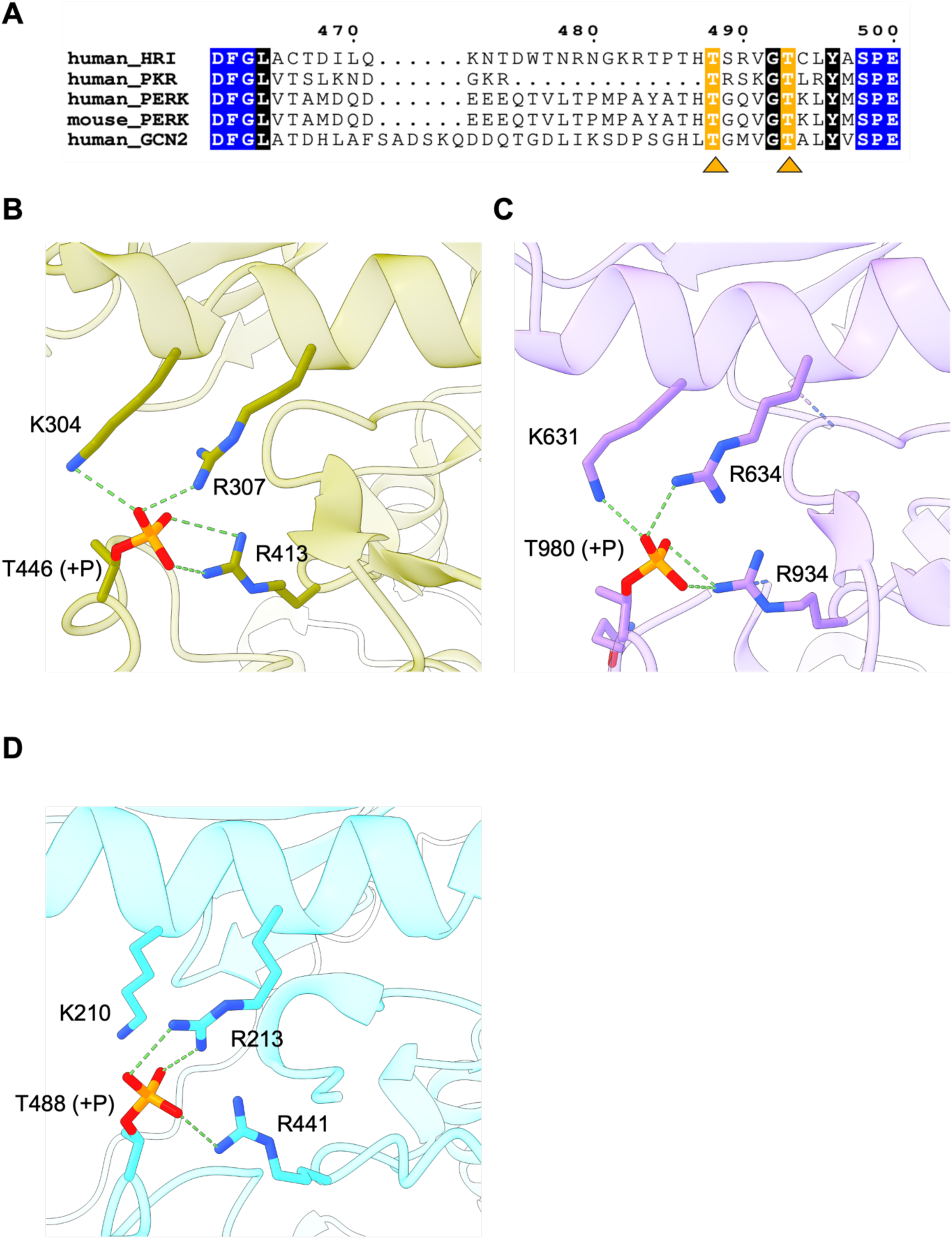
Structural comparison of autophosphorylation-mediated interactions in eIF2α kinases. (*A*) Multiple sequence alignment of the A-loop in eIF2α kinases. The alignment was performed using *Clustal Omega* and visualized with ESPript 3.0. Conserved residues are shown in black. Residues corresponding to Thr488/493 are highlighted in orange. The A-loop motif is shown in blue. UniProt accession numbers: PKR (P19525), PERK (Q9NZJ5 for human, Q9Z2B5 for mouse), GCN2 (Q9P2K8). **(*B–D*)** Representative interactions between phosphate groups and neighboring basic residues in **(*B*)** hPKR (PDB: 2A1A), **(*C*)** mPERK (PDB: 3QD2), and **(*D*)** HRI. PKR, PERK, and HRI are shown in khaki, purple, and cyan, respectively. Hydrogen bonds are indicated by green dotted lines.

### The regulatory mechanism of heme-induced inhibition of kinase activity

Previous studies have shown that the activation of HRI is regulated by its interaction with hemin, mediated by both the N-terminal and kinase domains (*7*,*19*,*22*,*32*). In particular, His119/120 and Cys410 in mouse HRI have been proposed as potential axial ligands for hemin. However, it should be noted that these experiments were primarily conducted using the phosphorylated form of HRI. We therefore examined hemin-induced regulation in dephosphorylated HRI to determine the domain primarily responsible for this effect.

First, the heme-binding capacity of HRI was determined through a hemin titration assay. Dephosphorylated HRIΔCC exhibited a specific absorbance peak at 419 nm upon heme titration (Fig. 10A), and dephosphorylated HRI-KD showed a peak at 415 nm (Fig. 10B), indicating that both proteins retain heme-binding capacity. We next evaluated kinase activity in the presence of hemin. The phosphorylation level of eIF2α decreased in a hemin concentration-dependent manner in both (-P)-HRIΔCC and (-P)-HRI-KD (Fig. 11A and B). Hemin significantly inhibited (-P)-HRIΔCC activity at concentrations of 20 and 40 μM after a 30-min reaction (Fig. 11C). Hemin also markedly inhibited the kinase activity of (-P)-HRI- KD at concentrations of 20 and 40 μM after a 120-min reaction, and the degree of inhibition was similar between (-P)-HRIΔCC and (-P)-HRI-KD. Moreover, hemin inhibited the autophosphorylation-induced band shifts of HRI in a concentration-dependent manner in both (-P)-HRIΔCC and (-P)-HRI-KD (Fig. 11D and E). Therefore, heme-induced deactivation occurs similarly in (-P)-HRIΔCC and (-P)-HRI-KD.

**Fig. 10.**
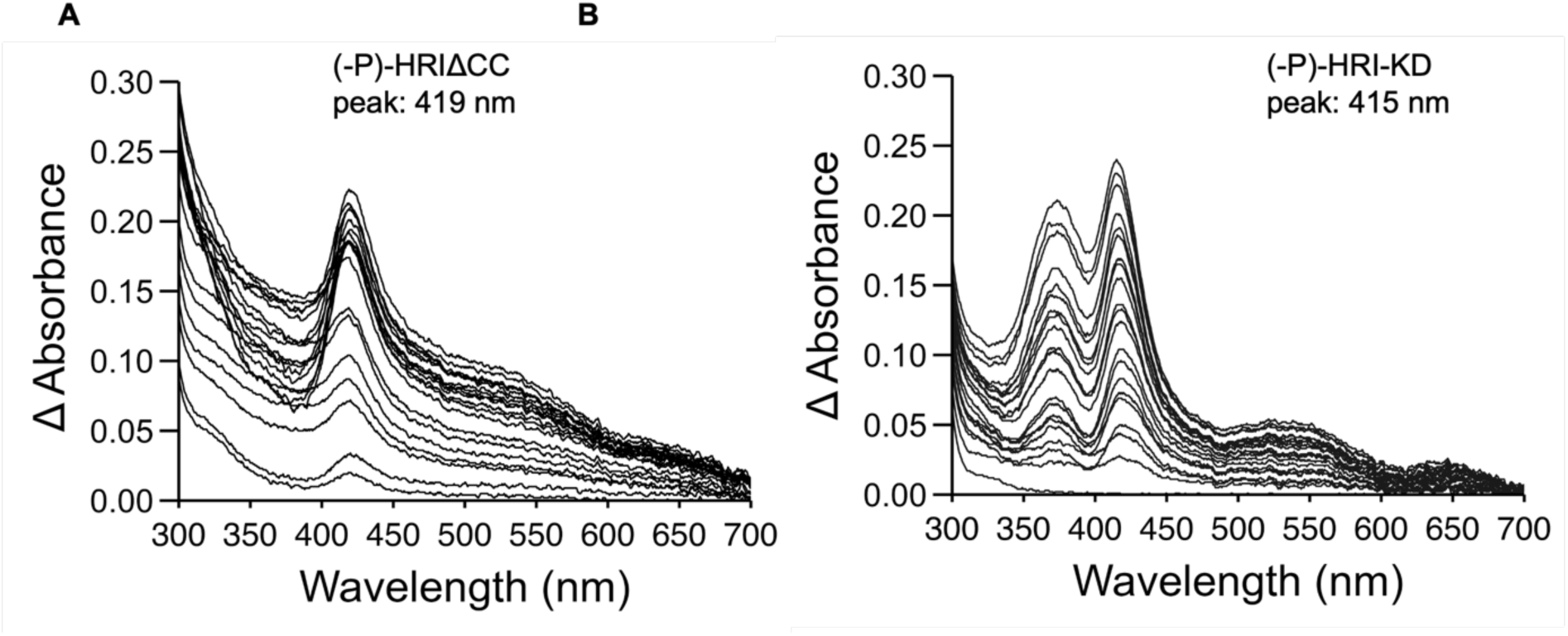
Heme-binding capacity of dephosphorylated HRI with or without the N- terminal domain. (*A, B*) Optical absorption spectra of **(*A*)** (-P)-HRIΔCC and **(*B*)** (-P)-HRI-KD following titration with hemin (2 µL increments). Absorption peaks were observed at 419 nm for (-P)-HRIΔCC and 415 nm for (-P)-HRI-KD.

**Fig. 11.**
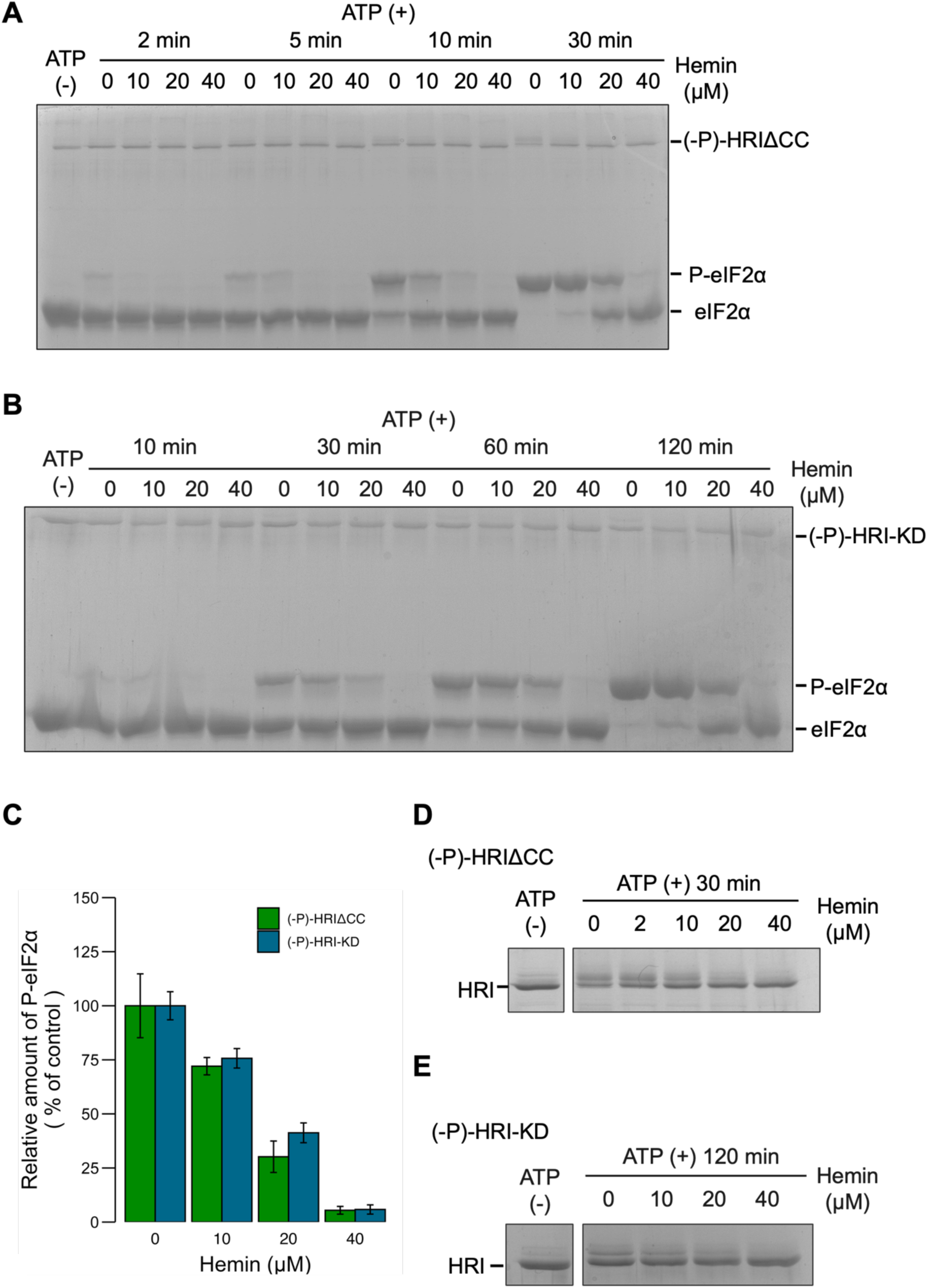
Heme-induced inhibition of kinase activity in dephosphorylated HRI with or without the N-terminal domain. (*A, B*) Kinase activity of **(*A*)** (-P)-HRIΔCC and **(*B*)** (-P)-HRI-KD in the presence of hemin at 0, 10, 20, and 40 µM. **(*C*)** Quantification of kinase activity based on **(*A*)** and **(*B*)**. Phosphorylated eIF2α levels at 30 min for (-P)-HRIΔCC and at 120 min for (-P)-HRI-KD (both at 0 µM hemin) were used as controls. Data represent the mean ± standard error of three independent experiments. **(*D, E*)** Autophosphorylation of **(*D*)** (-P)-HRIΔCC and **(*E*) (**-P)-HRI-KD under the same hemin concentrations.

To examine the function of the putative heme axial ligand in the kinase domain, we generated a dephosphorylated HRI-KD mutant in which Cys411 was replaced with Ser. We then evaluated kinase activity and heme-binding capacity (Fig. S2). As a result, eIF2α phosphorylation was still observed even under 40 μM heme treatment, although HRI-KD activity was significantly reduced under the same conditions (Figs. 11B, C, and S2A). This suggests that the Cys411 mutation alleviates hemin-induced inhibition. In addition, the mutant exhibited a specific peak at a wavelength of 417 nm (Fig. S2B), closely resembling that of (-P)-HRI-KD, indicating that the mutant still retains heme-binding capacity.

## Discussion

HRI primarily regulates hemoglobin translation in response to endogenous heme levels; however, the mechanisms of autophosphorylation-induced activation and heme sensing remain unclear. Our study demonstrates that HRI forms a dimer in solution in an N-terminal domain-dependent manner, regardless of its autophosphorylation state (Figs. 2B and 3B). A recent report showed that full-length HRI dimerizes through its C-terminal coiled-coil domain (*22*). However, our recombinant HRI lacking the C-terminal coiled-coil domain remained a stable dimer through the N-terminal domain. Together, these findings suggest that full-length HRI forms a stable dimer in solution through both its N-terminal domain and its C-terminal coiled-coil domain. The structure predicted by AlphaFold 3 indicated dimerization of the N-terminal domains (Fig. 12A). The predicted dimer interface consisted of hydrophobic residues, including Leu68, Leu69, Leu72, Leu76, and Val79, which are likely to form hydrophobic interactions (Fig. 12B). Consistent with this, previous reports indicated that these residues were not autophosphorylation sites (*19*). Therefore, these hydrophobic interactions are likely responsible for maintaining N-terminal domain-dependent dimerization regardless of the phosphorylation state. Our results further suggest that the N-terminal domain enhances enzymatic activity (Fig. 2C–F), consistent with previous findings (*32*). However, the monomeric kinase domain retained autophosphorylation activity despite lacking the N-terminal domain (Fig. 3C–H). In addition, earlier studies reported that the C-terminal coiled-coil domain is not required for autophosphorylation activity (*22*). Together, these results suggest that HRI dimerization via either the N-terminal domain or the C-terminal coiled-coil domain is not a prerequisite for autophosphorylation. HRI contains many phosphorylation sites, but how these sites undergo autophosphorylation remains unclear. For comparison, the kinase domain of PKR forms two distinct dimers: a back-to-back dimer mediated by the N-lobes and a front-to-front dimer formed through domain swapping (*33*, *34*). The latter conformation is considered relevant to autophosphorylation, as it promotes a catalytically favorable configuration. By analogy, HRI may also adopt alternative oligomeric states to facilitate autophosphorylation.

**Fig. 12.**
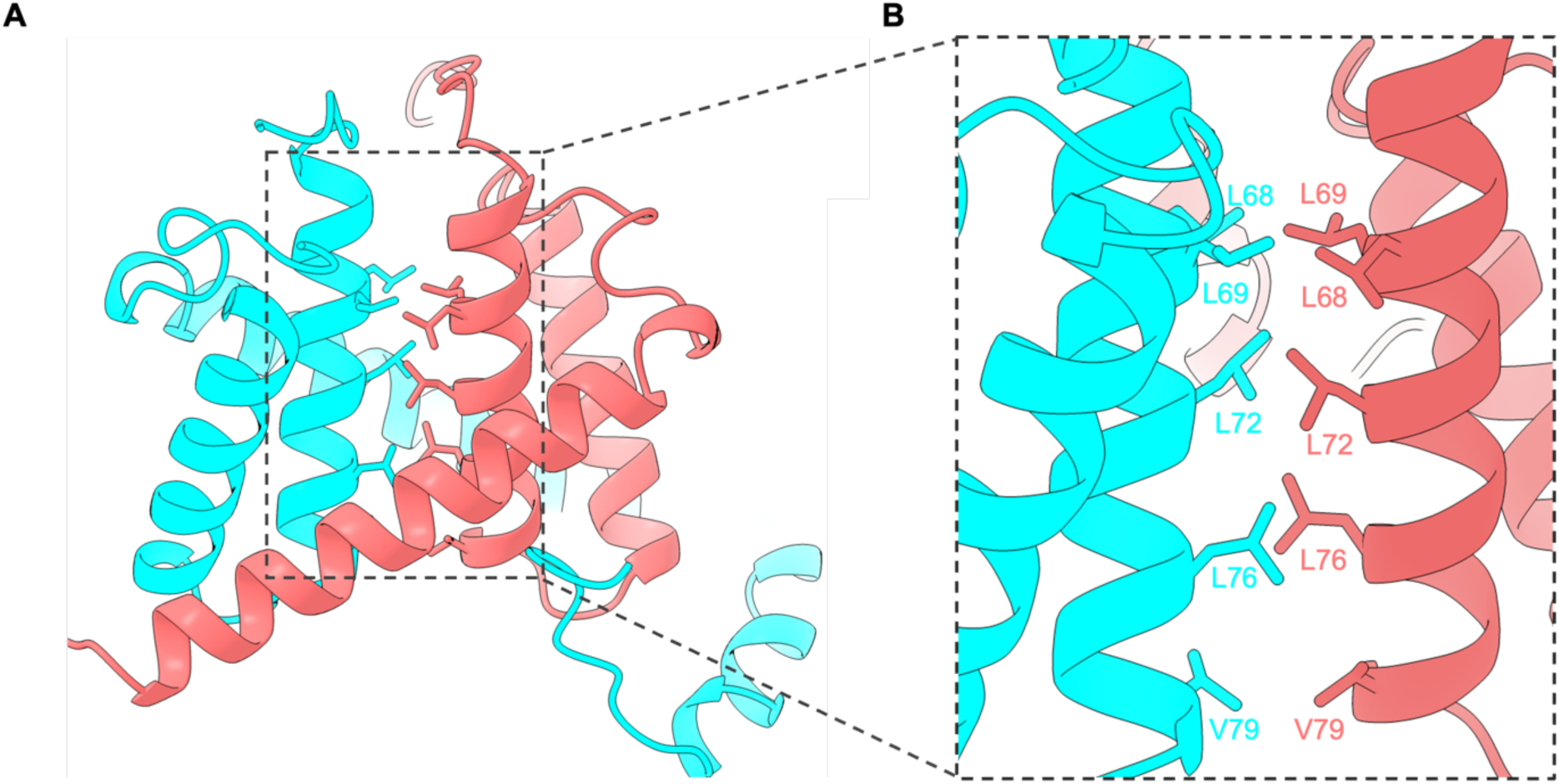
Predicted N-terminal domain-mediated dimerization. (*A*) Predicted dimer model of the N-terminal domain generated by AlphaFold 3. The two N-terminal domains are shown in cyan and red. **(*B*)** Hydrophobic interactions observed at the dimer interface.

We also examined the mechanism by which autophosphorylation regulates HRI activity. Given that dephosphorylated HRI was unable to interact with eIF2α (Fig. 4), our findings suggest that autophosphorylation is essential for eIF2α binding. Notably, dephosphorylated HRI retained kinase activity in the presence of ATP, showing levels comparable to phosphorylated HRI (Fig. 3C–H), and it completely phosphorylated eIF2α even while autophosphorylation was ongoing. These results suggest the existence of a minimal set of autophosphorylation sites required for activation. In particular, autophosphorylation at Thr488 and Thr493 appears to enhance eIF2α binding and kinase activity by enabling their phosphate groups to interact with nearby basic residues (Figs. 6–8). Our results further suggest that phosphorylation of Thr488 and Thr493 induces a conformational change in the A-loop that favors eIF2α interaction. The crystal structures of hPKR and mPERK also show similar interactions (Fig. 9). Furthermore, Thr446/Thr451 in hPKR and Thr882/Thr887 in yeast GCN2, corresponding to Thr488/Thr493 in HRI, have been reported to play essential roles in kinase activity (*35*). Taken together, these findings suggest that autophosphorylation-induced conformational changes in the A-loop represent a conserved mechanism among eIF2α kinases. Nevertheless, replacing Thr488 and Thr493 with alanine did not completely abolish eIF2α binding, unlike in the dephosphorylated form (Figs. 4 and 7A–D), suggesting that other autophosphorylation sites also contribute to this interaction. Deletion of the kinase insert loop markedly reduced eIF2α binding to levels comparable to that of dephosphorylated HRI (Figs. 4A, C, and 6E, F). These results highlight the essential role of the kinase insert loop in eIF2α recognition. In mHRI, the loop contains 17 autophosphorylation sites (*19*), suggesting that phosphorylation can dynamically alter its conformation to enhance affinity for eIF2α. Notably, the kinase insert loop is not conserved among other eIF2α kinases. Therefore, HRI employs a unique mechanism for eIF2α recognition, relying on both the A-loop and the kinase insert loop to regulate binding affinity. Further studies are needed to identify additional autophosphorylation sites that modulate the catalytic activity and eIF2α interaction.

Our study indicates that the heme-induced inhibition of kinase activity occurs primarily within the kinase domain of dephosphorylated HRI. In addition, autophosphorylation was also inhibited by heme treatment (Figs. 10 and 11). A recent preprint in bioRxiv reported that heme binding induces a global conformational change in dephosphorylated HRI (*36*). Hydrogen-deuterium exchange mass spectrometry revealed altered solvent exchange rates across many residues under heme treatment, with notably reduced exchange rates in residues within the A-loop (*36*). These findings suggest that heme induces dynamic structural changes in dephosphorylated HRI, which may cause the A-loop to become shielded by other regions. We also confirmed that the dephosphorylated HRI-heme complex adopts distinct oligomeric states compared with apo HRI (data not shown). Taken together with previous reports, our results suggest that heme-induced structural changes may contribute to autophosphorylation inhibition, particularly within the kinase domain. Previous research showed that the heme-induced inhibition of kinase activity is more pronounced in phosphorylated HRI containing both the N-terminal and kinase domains (*7*). In contrast, catalytic inhibition in dephosphorylated HRI was similar regardless of the presence of the N-terminal domain (Fig. 11C). In the dephosphorylated state, heme inhibited autophosphorylation (Fig. 10D, E), whereas this regulation appears not to occur in the already-phosphorylated form. This suggests the existence of other regulatory mechanisms depending on the phosphorylated form. In summary, the different roles of the N-terminal domain in phosphorylated and dephosphorylated forms with respect to heme-induced activity inhibition may reflect distinct regulatory mechanisms. Previous studies suggested that Cys410 in the kinase domain of mHRI coordinates with heme (*7*). Consistent with this, our results showed that the heme-induced inhibition was similarly suppressed by mutating hHRI Cys411 corresponding to mHRI Cys411 to Ser in the kinase domain of dephosphorylated hHRI, as previously observed in mHRI. Nevertheless, the mutant retained the heme-binding capacity, similar to WT HRI-KD (Figs. 10 and S1B). Besides the CP motif, other amino acids may also contribute to heme interaction. Further studies, especially structural analyses, are required to identify the residues involved in heme binding. Our study provides mechanistic insights into the regulation of HRI activity by autophosphorylation and heme binding. These findings advance our understanding of the regulatory mechanisms governing HRI.

## Abbreviations

A-loop: activation loop
CP motif: Cys-Pro motif
eIF2α: eukaryotic translation initiation factor 2 subunit α
GCN2: general control nondepressible 2
HRI: heme-regulated inhibitor
ISR: integrated stress response
KD: kinase domain
MW: molecular weight
PERK: protein kinase R-like endoplasmic reticulum kinase
PKR: protein kinase R
SEC: size exclusion chromatography
Sw: sedimentation coefficient
WT: wild type

## Data availability

The data that support the findings of this study are available from the corresponding author upon reasonable request.

## Authors’ contributions

H.Y. and S.Y.P. conceived the study. T.O., K.M., E.O., and H.Y. designed and performed the experiments. T.O. and K.I. drafted the manuscript. H.Y., S.Y.P., and K.I. supervised the experiments, and reviewed and edited the manuscript. All authors approved the manuscript.

## Acknowledgement

We thank Dr. Kenichi Kamata (formerly at Yokohama City University, now at Nihon University) for his assistance with the instrument.

## Supporting information

**Fig. S1.**
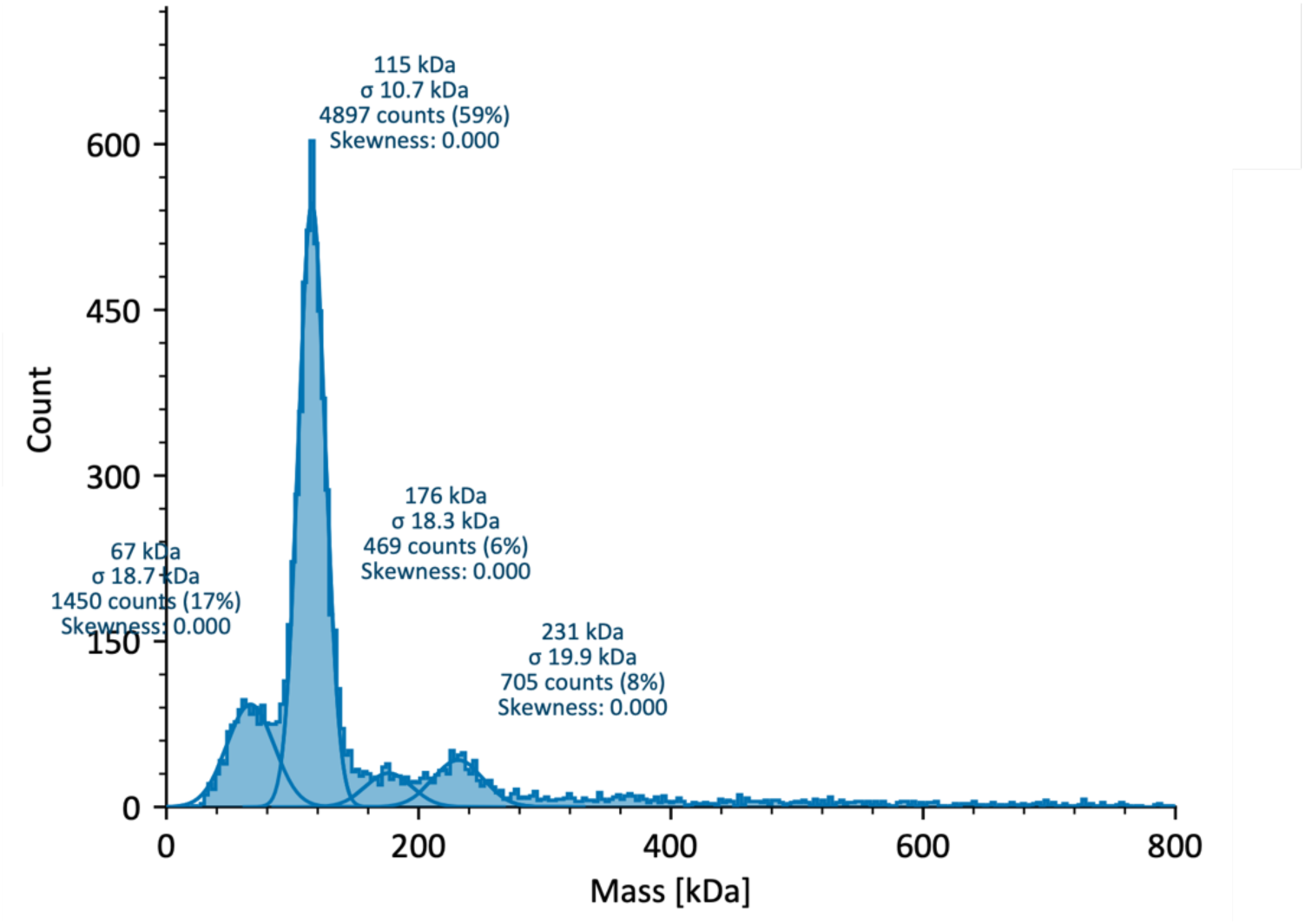
**Mass photometry of (-P)-HRI to assess its oligomeric state in solution.**

**Fig. S2.**
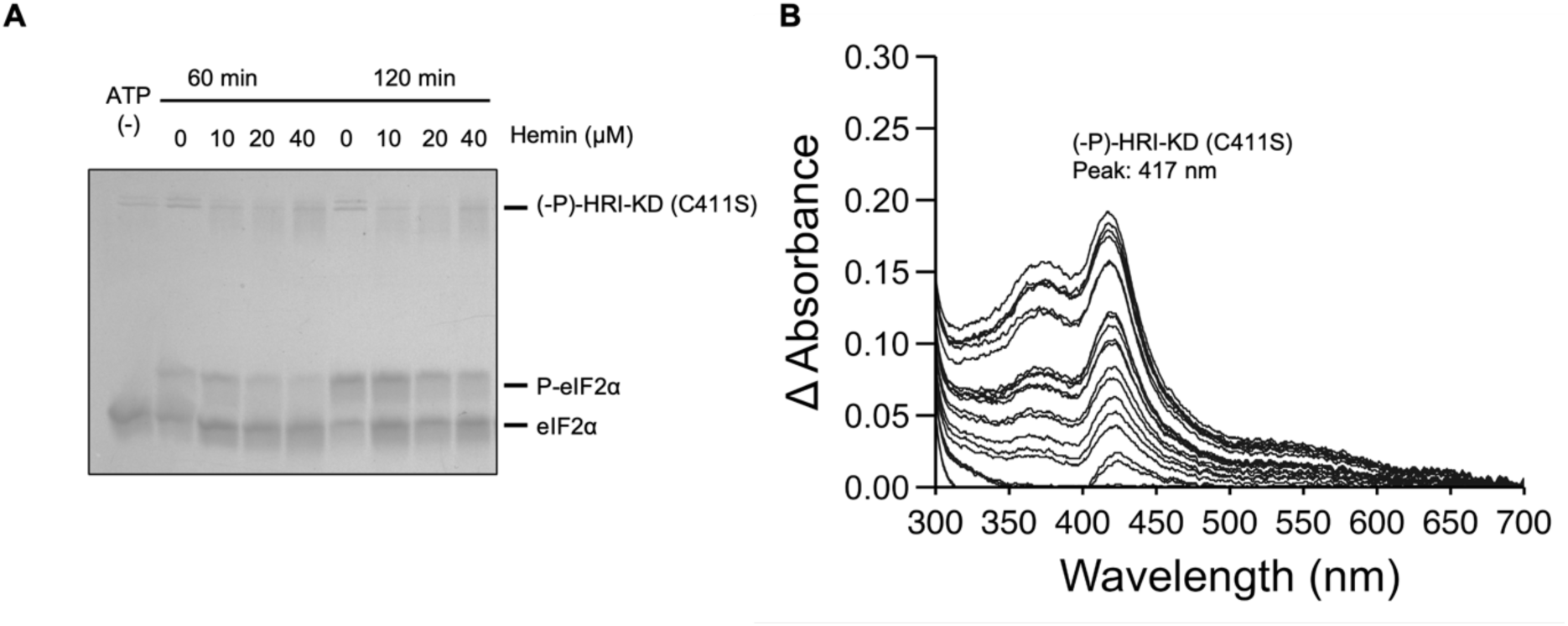
Hemin-induced kinase inhibition and heme-binding capacity of (-P)- HRI-KD (C411S). **(A)** Kinase activity of (-P)-HRI-KD (C411S) in the presence of hemin. **(*B*)** Optical absorption spectra of (-P)-HRI-KD (C411S) following hemin titration (2 µL increments). The absorption peak was observed at 417 nm.

